## Supplementary Figures for "A male-drive female-sterile system for the self-limited control of the malaria mosquito *Anopheles gambiae*": Strampelli et al SUPPLEMENTARY FIGURES.docx

**Supplementary Figure 1**

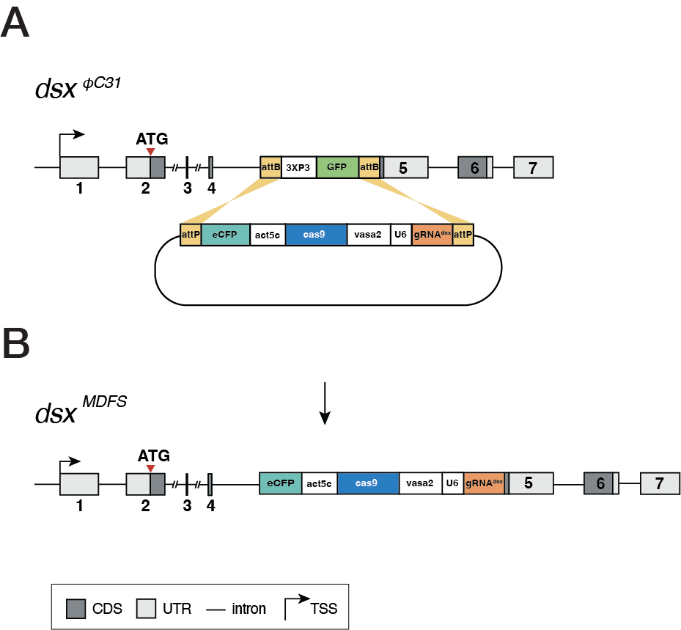

**Supplementary Figure 1.** Recombinase-mediated cassette exchange was used to swap out the *dsx*^φC31^ allele with the MDFS construct. (**A**) This involved using the catalytic activity of the φC31 integrase to replace the *dsx*^φC31^ allele, which contains the 3xP3::GFP transcription unit surrounded by *attB* sites, with the MDFS construct. The MDFS construct consists of an *actin5c::eCFP* fluorescent marker, a *Cas9* driven by the germline *vasa2* promoter and a *U6::gRNA^dsxF^* cassette targeting *dsx* at the intron 4-exon 5 boundary, all surrounded by two *attP* sites. (**B**) The schematic depicts the MDFS construct within the *dsx* gene post-integration. The insertion of the construct hinders the production of functional *dsxF* transcript, leaving *dsxM* unaffected. Non-coding regions (UTR) are shaded in light grey, coding regions (CDS) are shaded in dark grey, and black lines indicate introns and are not in scale. The arrow denotes the transcription start sites (TSSs/promoter) of the *dsx* gene.

**Supplementary Table 1**

| **G1** | **MDFS**  **(CFP+)** | ***attP* docking site (GFP+)** | **Sex assigned in pupae** | **Sex assigned in adults** |
| --- | --- | --- | --- | --- |
| 1 | ✔ | ✔ | **♂** | **♂** |
| 2 | ✔ | ✔ | **♂** | **♂** |
| 3 | ✔ | ✔ | **♂** | **♂** |
| 4 | ✔ | ✔ | **♂** | **♂** |
| 5 | ✔ | ✔ | **♂** | **♂** |
| 6 | ✔ | X | **♂** | **♀** (intersex) |
| 7 | ✔ | X | **♂** | **♀** (intersex) |
| 8 | ✔ | ✔ | **♂** | **♀** (intersex) |

**Supplementary Table 1.** G1 transgenics obtained upon injections of the MDFS construct in the *attP* docking site embryos. The docking strain used for the embryo injections mainly consisted of homozygous individuals for the *dsx*^φC31^ allele. G1 transgenics were expected to be CFP^+^ if they contained only the MDFS allele or both CFP^+^ and GFP^+^ if they contained the MDFS allele and the *dsx*^φC31^ allele. G1 male number 2, highlighted in light blue, was arbitrarily selected to establish the MDFS strain.

**Supplementary Figure 2**

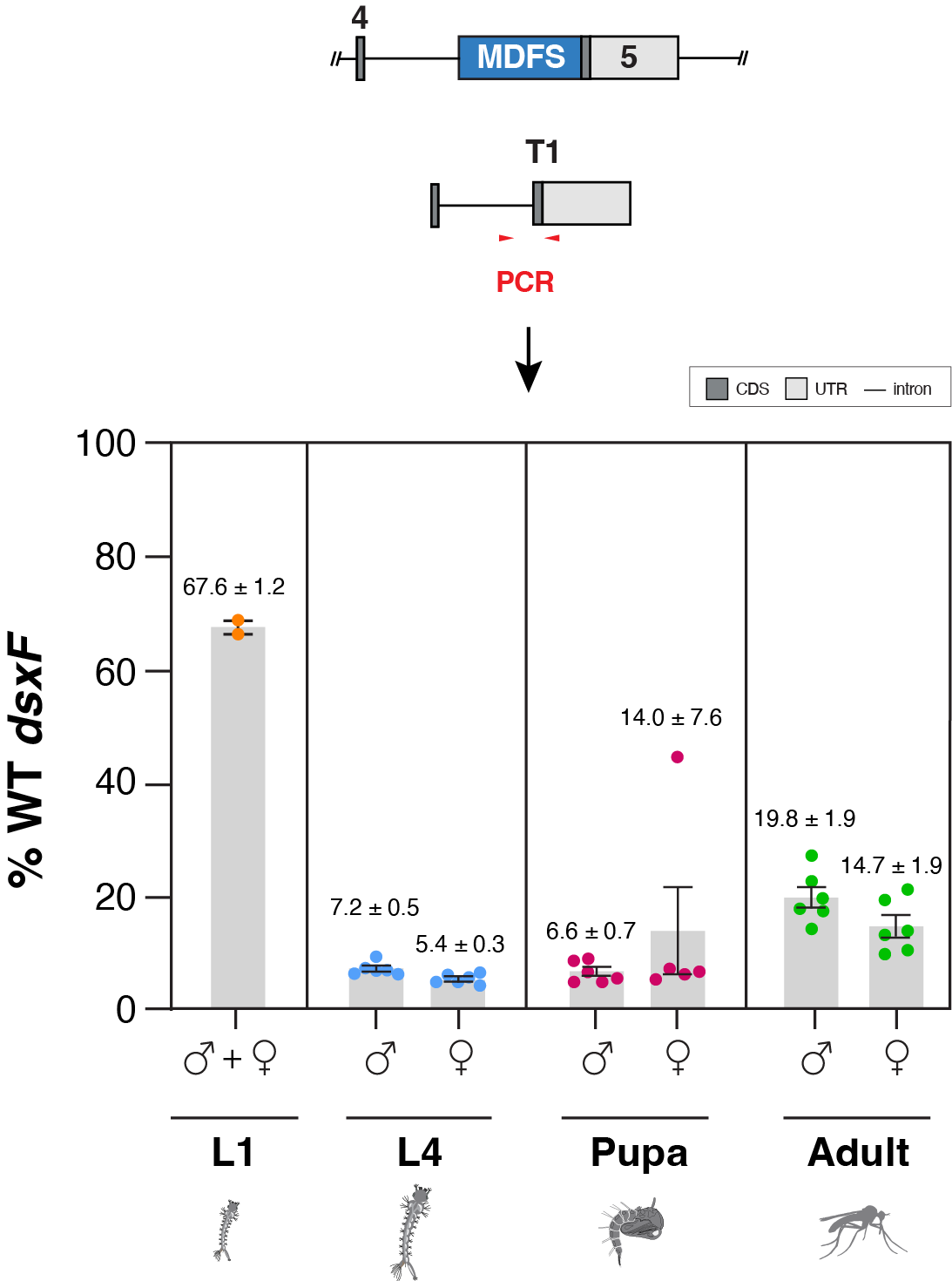

**Supplementary Figure 2.** Frequency of wild-type alleles at the *dsxF* target site in MDFS individuals at different developmental stages. We performed a pooled amplicon sequencing analysis at the *dsxF* target site to quantify the level of disruption at different life stages of male and female MDFS individuals. The wild-type (WT) dsxF frequency refers to the allele that does not contain the MDFS transgene (the PCR symbol is placed at the top of the diagram to symbolize the region amplified). L1 samples were two pools of >100 larvae, while each dot represents a single individual for the other stages. Since L4 male and female larvae are phenotypically indistinguishable, and MDFS female pupae are phenotypically male-like, to separate males and females at these two developmental stages, we crossed MDFS males to females homozygous for a mCherry marker under the male-specific *β2* promoter. The male progeny of this cross expressed mCherry in the testes, and the female progeny did not, which allowed the two sexes to be separated for the analysis. Horizontal bars indicate the mean and the s.e.m..

**Supplementary Figure 3**

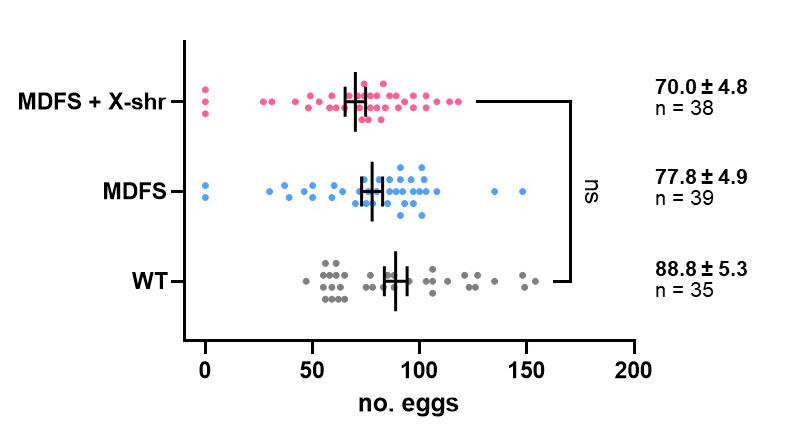

**A**

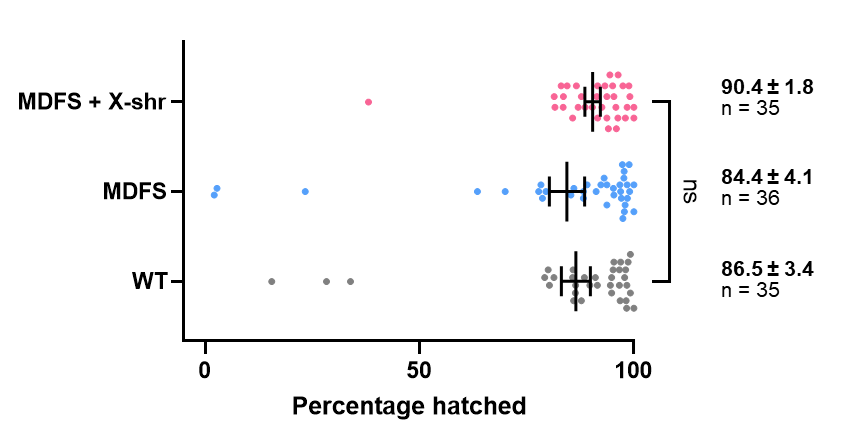

**B**

**Supplementary Figure 3.** Fertility assay of MDFS and MDFS + X-shredder males. **(A)** The egg output and **(B)** hatching rate (larvae/eggs) were measured and compared to those of wild-type (WT) controls. No significant differences (‘ns’) were found among these genotypes (P>0.05; Kruskal-Wallis test adjusted for multiple comparisons). Vertical bars indicate the mean and the s.e.m..

**Supplementary Figure 4**

**A**

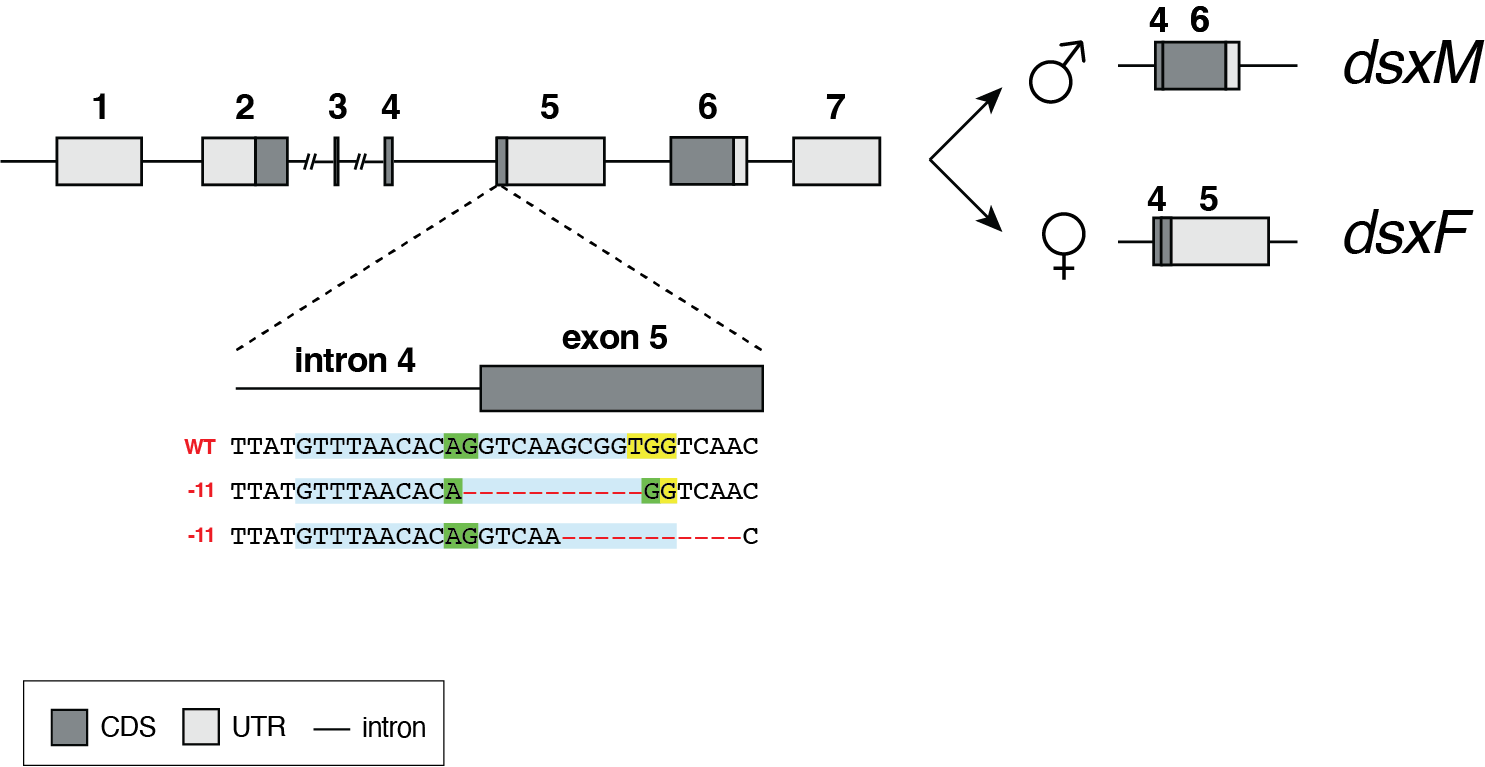

**B**

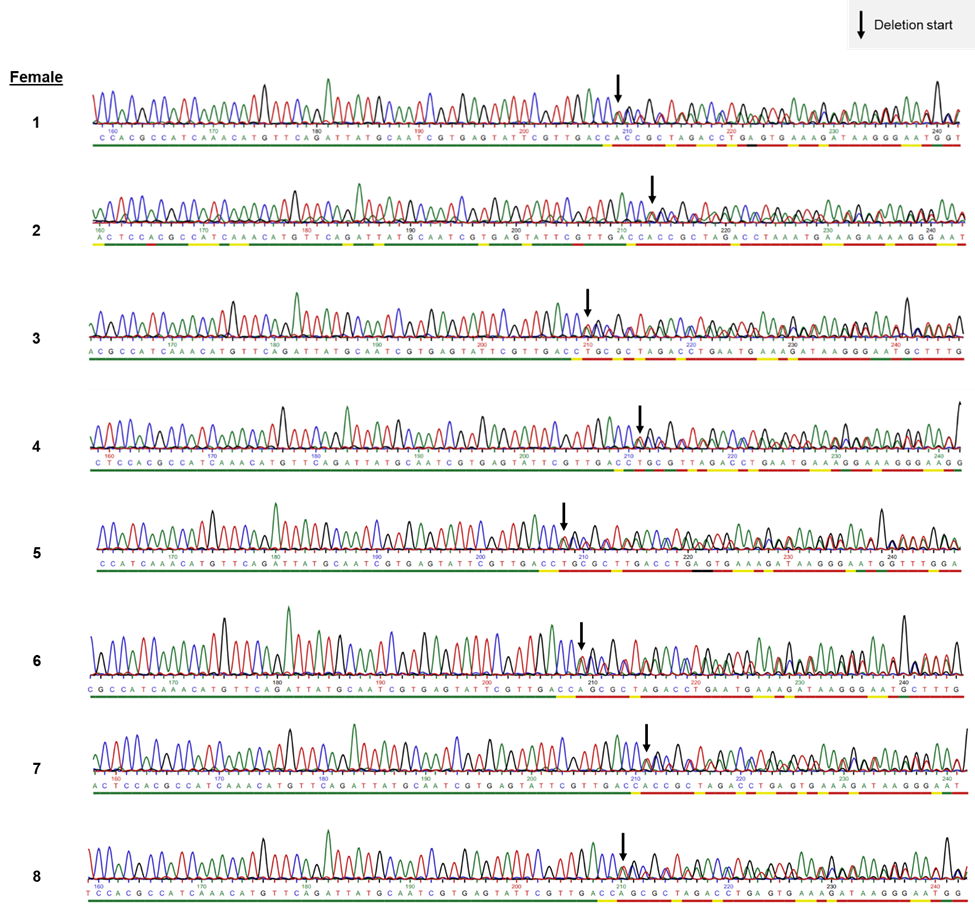

**Supplementary Figure 4.** Deletions identified in the eight mosaic females that did not inherit the MDFS allele (CFP-). **(A)** The 11 bp deletion aligned against the wildtype allele, which could be portrayed in one of two ways because it resulted from microhomology-mediated end joining (MMEJ) repair. The AG sequence of the splice acceptor site is retained and highlighted in green, and the PAM is highlighted in yellow. **(B)** The electropherograms showing the Sanger sequences of the *dsxF* target site in the females analyzed, where the 11-bp deletion begins at the transition from one to double peaks and is indicated by the black arrow. Note that the sequencing was performed with a reverse primer with respect to the *dsxF* sequence, so the “GTTGACC” sequence that is 5’ of the deletion in (A) is the reverse complement of the “GGTCAAC” sequence that is 3’ of the deletion in (B).
